# Frontal engagement in perceptual integration under low subjective visibility

**DOI:** 10.1101/2024.03.12.584542

**Authors:** Jisub Bae, Koeun Jung, Oliver James, Satoru Suzuki, Yee Joon Kim

**Affiliations:** Center for Cognition and Sociality, Institute for Basic Science (IBS), Daejeon, South Korea; Institute of Social Science, Chungnam National University, Daejeon, South Korea; Department of Psychology, Northwestern University, Evanston, IL 60208, USA

**Author notes:** Corresponding author, Correspondence: Yee Joon Kim, PhD, Center for Cognition and Sociality, Institution for Basic Science, 55 Expo-ro, Doryong-dong, Yuseong-gu, Daejeon, 34126, South Korea. These authors contributed equally: Jisub Bae, Koeun Jung.

**Keywords:** Kanizsa illusory contour, Perceptual integration, Consciousness, Multivariate pattern analysis, Temporal generalization method

## Abstract

We investigated how spatiotemporal neural dynamics underlying perceptual integration changed with the degree of conscious access to a set of backward-masked pacman-shaped inducers that generated the percept of an illusory triangle. We kept the stimulus parameters at a fixed near-threshold level throughout the experiment and recorded electroencephalography from participants who reported the orientation and subjective visibility of the illusory triangle on each trial. Our multivariate pattern analysis revealed that posterior and central regions initially used dynamic neural code and later switched to stable neural code. The transition from dynamic to stable neural code in posterior region occurred increasingly later and eventually disappeared with decreasing conscious access. Anterior region primarily used stable neural code which waned with decreasing conscious access, but increased at below-median visibility and remained even when stimulus awareness was minimal. These results demonstrate differential spatiotemporal neural dynamics underlying perceptual integration depending on conscious access and emphasize a unique role of anterior region in processing integrated shape information especially under low subjective visibility.

## Introduction

Perceptual integration, the process of combining fragmented sensory information into a cohesive whole, is indispensable for detecting and identifying objects in natural environments where occlusion is commonplace. While many aspects of perceptual integration have been investigated, the relationship between perceptual integration and conscious access has been the subject of ongoing debate, particularly with regard to the possibility of perceptual integration occurring without conscious access^1–15^. In most studies using attentional blink or masked priming paradigm, conscious access was determined based solely on the level of target detection performance without any report of target visibility. Thus, it has been implicitly assumed that there is no conscious access when participants perform the target detection task at chance level. However, recent research has shown that a stimulus close to threshold yields a mixture of high and low conscious access trials, producing a burst in the trial-to-trial variability of conscious access^16^. Therefore, it remains unclear whether perceptual integration is independent of conscious access.

Here, we used the Kanizsa illusion to investigate how conscious access to the physically identical backward-masked Kanizsa figures varied, and how this variation in the degree of conscious access altered the spatiotemporal neural dynamics underlying the integrated percepts of Kanizsa illusory contours. Especially, we focused on the near-threshold level of conscious access, where behavioral performance was near or at chance level and Kanizsa visibility was in a nebulous in-between state of visible and invisible. To capture the spatiotemporal neural dynamics of perceptual integration across different levels of conscious access while keeping the stimulus parameters constant, we first optimized the contrast and temporal parameters of the backward mask to ensure that the visibility of the target Kanizsa stimuli substantially varied from trial to trial purely due to changes in internal brain states. However, when there is no physical change in stimulus conditions across trials, target identification accuracy alone is not sufficiently informative of the degree of conscious access to the target stimulus as the stimuli from incorrect trials may still have been clearly seen or the stimuli from correct trials may have been unseen at all. We thus had participants report not only the identity but also the subjective visibility of Kanizsa stimuli, indicating whether a Kanizsa triangle was not detected at all, detected with a vague sense of its orientation, or detected with a clear sense of its orientation (see Methods). This design provided a unique opportunity to investigate brain responses as a function of conscious access without confounding effects from varying stimulus strengths, unlike the previous related studies that manipulated stimulus parameters^5,17,18^.

Further, we evaluated the spatiotemporal neural dynamics underlying perceptual integration to determine whether integrated percepts are reflected in a stable and/or a time-changing activity pattern^19,20^, by applying multivariate pattern analysis in conjunction with high-temporal resolution electroencephalography (EEG) across space and time. Stable neural code refers to a situation where a content-specific information is reflected in sustained activity patterns over time, so that the pattern at one time can also reconstruct the same information at another time. In contrast, dynamic neural code refers to a situation where a content-specific information is reflected in dynamically changing activity pattern that can only be reconstructed at the time point when the decoder is built. As the roles of spatiotemporal profiles of conscious and unconscious processes have been extensively debated^6,21,22^, we applied this method to the anterior, central, and posterior groups of electrodes separately and quantified the spatiotemporal dynamics with a dynamic/stable index^20^ to determine whether the temporal dynamics of perceptual integration are dynamic or stable in different brain regions and, furthermore, how the spatiotemporal dynamics are affected by the degree of conscious access.

The full spatiotemporal dynamics revealed a sequence of distinct brain activation patterns engaged in perceptual integration in different brain regions. Both posterior and central regions initially used dynamic neural code and later switched to stable neural code. This transition from dynamic neural code to stable neural code in posterior region occurred increasingly later and eventually disappeared as Kanizsa visibility decreased. However, anterior region mainly used stable neural code that supported the significant decoding of integrated percepts. Especially when behavioral performance was at chance level and Kanizsa visibility was near median anterior region primarily used stable neural code during the almost entire period from 300 ms to 1000 ms after the stimulus onset. The stable neural code in anterior region still remained even when Kanizsa visibility was minimal. Taken together, these results demonstrate the differential spatiotemporal neural dynamics underlying perceptual integration depending on subjective visibility of constant stimuli and emphasize the role of frontal region in perceptual integration especially at low subjective visibility.

## Results

We analyzed scalp EEG signals from 20 observers while they viewed either a Kanizsa target or a control target (Figure 1a). Each target was presented for 20 ms, followed (after a 40 ms blank interval) by a 20 ms mask (Figure 1b). Targets were presented in a random sequence and could be either one of the four Kanizsa stimuli (with the emergent triangle pointing in upper-right, upper-left, lower-left, and lower-right directions) or a control stimulus (Figure 1a). Observers were instructed to rate the visibility (0-7) of a Kanizsa triangle and to indicate the identity of the target stimulus (guessing if necessary). Identification of Kanizsa targets was significantly above chance on high-visibility trials (visibility = 4-7), but it was not significantly above chance on low-visibility trials (visibility = 0-3) (Figure 1c, p<0.0001, Bonferroni corrected).

**Figure 1.**
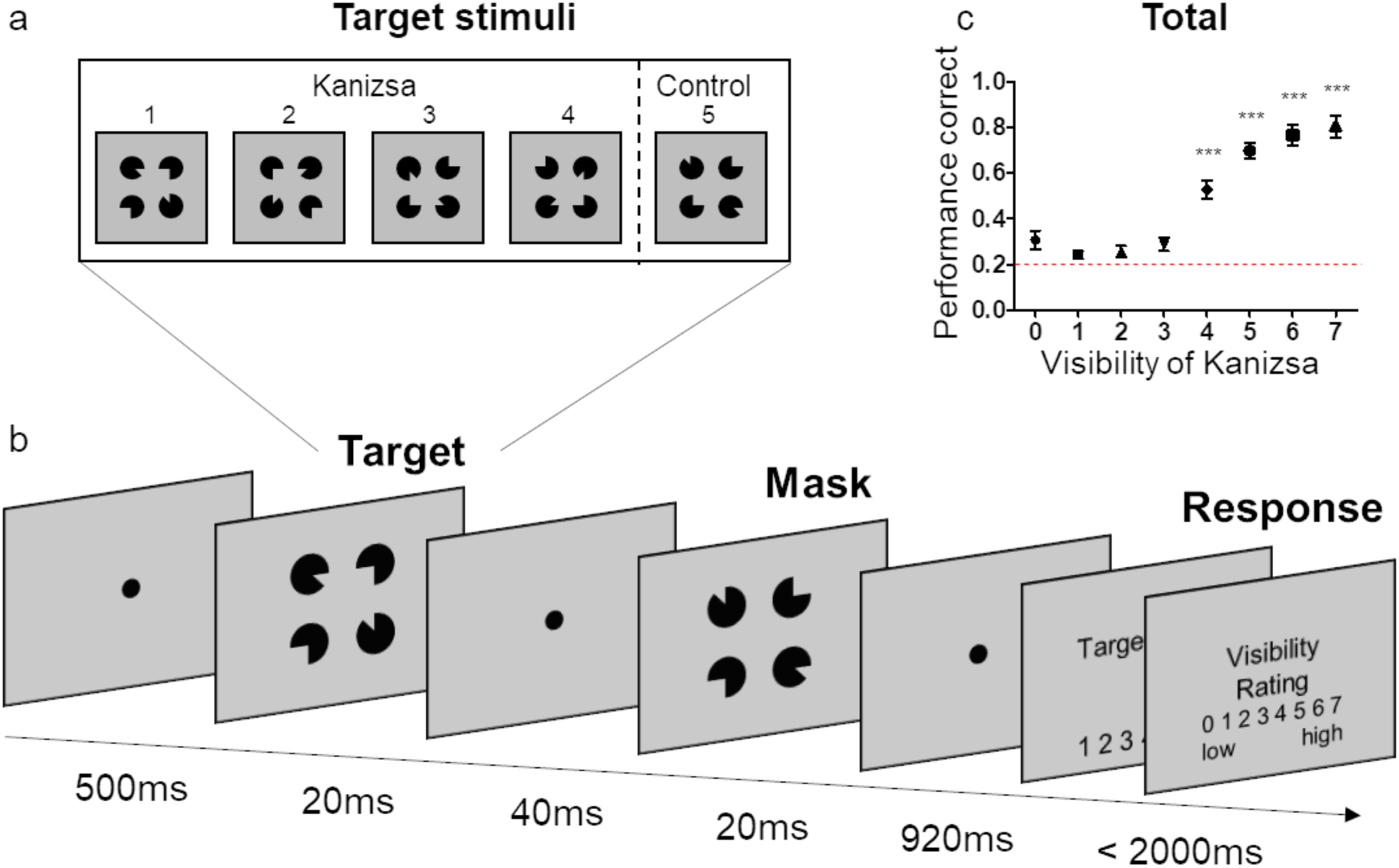
Stimuli and procedure for the behavioral experiment. **a.** The Kanizsa and control stimuli. **b.** Trial events. After a 500 ms fixation period, a target stimulus was presented for 20 ms, followed by a 20 ms mask after 40 ms inter-stimulus interval. During the subsequent response period, observers indicated the target they saw (1 to 5) and then rated the visibility (1 to 7) of a Kanizsa triangle. **c.** Target identification accuracy as a function of Kanizsa visibility. The red dashed line indicates chance level. Error bars indicate ±1 SEM. ***p<0.0001.

To establish a neural index of perceptual integration, we trained a linear discriminant analysis (LDA) classifier to categorize the trials as either Kanizsa or control using the EEG signals across electrodes as features. We calculated the ability to discriminate between the Kanizsa and control stimuli as the area under the receiver operating characteristic curve (AUC) computed from the true positive rate (TPR, the proportion of Kanizsa trials correctly categorized as Kanizsa) and false positive rate (FPR, the proportion of control trials incorrectly categorized as Kanizsa). We used the AUC as a measure of the accuracy of decoding the presence of perceptual integration.

### Frontal engagement in spatiotemporal dynamics of perceptual integration emerges during the shift from high to low visibility conditions

To systematically explore how neural dynamics underlying perceptual integration changed as a function of conscious access to a Kanizsa triangle using multivariate pattern analysis, we sorted Kanizsa trials in order from correct trials ranked by visibility rating to incorrect trials ranked by visibility rating. We repeated the same trial sorting procedure for each Kanizsa stimulus type. From these sorted trials, we selected the top 50 and bottom 50 trials of each of the four Kanizsa triangle types to create Dataset 1 (200 trials in total) and Dataset 4 (200 trials in total), respectively. As conscious access is an issue when task performance is at chance level according to previous studies^1–5^., we created Dataset 3 by extracting trials in which behavioral performance was around chance level of 0.2. As the average visibility rating of Dataset 3 was 3, which was just below the grand average visibility rating of 3.5, we created Dataset 2 by extracting trials that made the average visibility rating higher than the grand average visibility rating by 0.5. In this way, Dataset 2 and Dataset 3 mainly consisted of trials with ambiguous conscious access, such as correct trials with low visibility ratings or incorrect trials with high visibility ratings, and they were positioned between Dataset 1 and Dataset 4. Thus, trials were split into four datasets of equal number of trials in the decreasing order of conscious access: the maximal visibility dataset (Dataset 1, visibility *M* = 5.3, performance correct *M* = 98%), above-mean visibility dataset (Dataset 2, visibility *M* = 4, performance correct *M* = 60%), below-mean visibility dataset (Dataset 3, visibility *M* = 3, performance correct *M* = 16%), and minimal visibility dataset (Dataset 4, visibility *M* = 1.6, performance correct *M* = 8%). Figure 2a shows the behavioral performance on the Kanizsa stimuli and the average Kanizsa visibility rating for each dataset. Stimulus identification performance was no greater than chance for Dataset 3 and 4 (The 1^st^ column of Fig 2a). For the control dataset, we used all 200 trials of non-Kanizsa control stimuli. As expected, the Kanizsa visibility rating was low in response to the control stimuli (visibility, *M* = 1.3).

**Figure 2.**
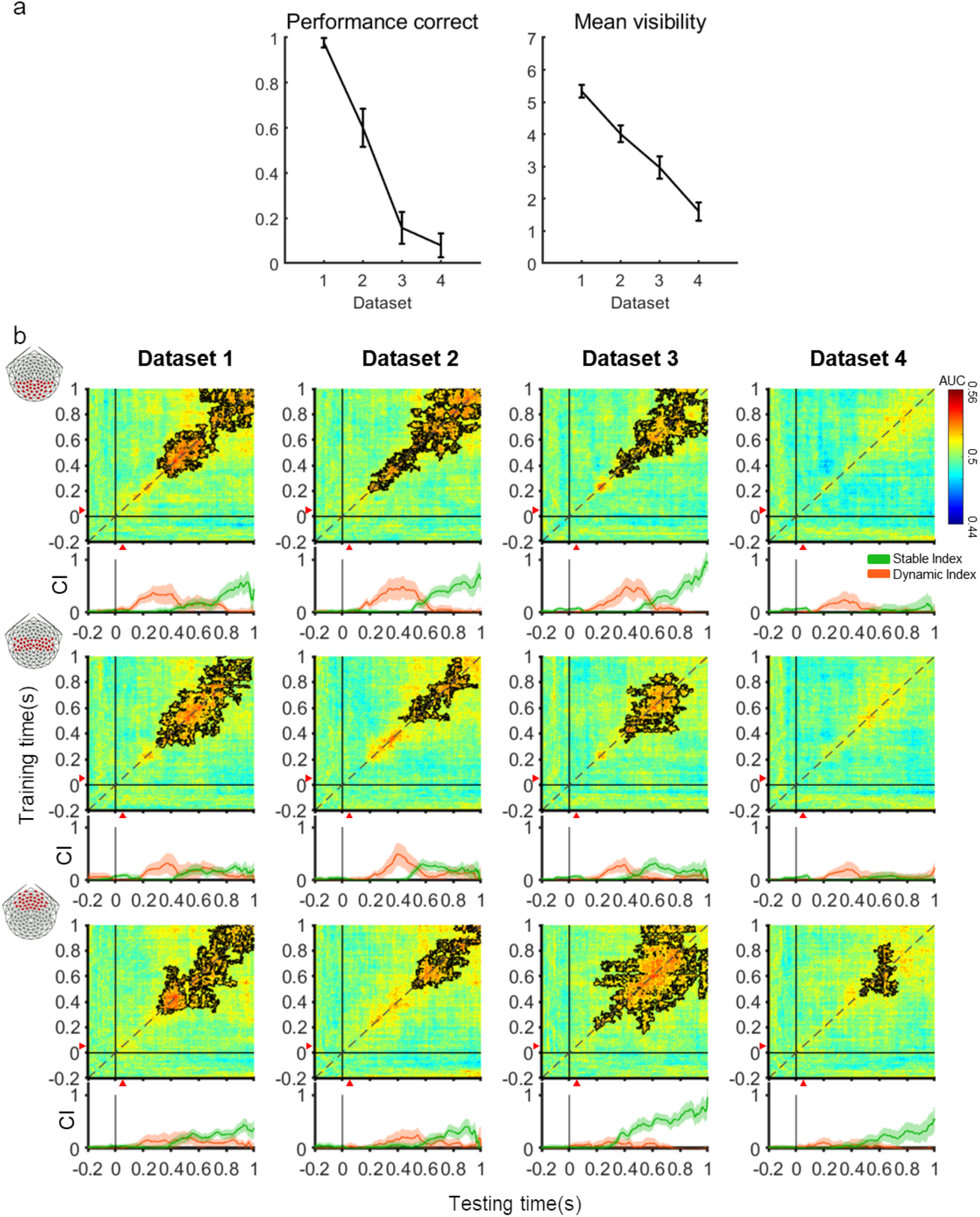
Spatiotemporal neural dynamics of perceptual integration as a function of subjective visibility. **a.** Performance correct and average Kanizsa visibility rating as a function of dataset. **b.** TG maps and dynamic/stable index of each dataset for the posterior, central, and anterior region (see electrode maps on the left). Colors represent the decoding accuracy (AUC). Black contours indicate clusters of significant decoding accuracies above chance (p < 0.05, based on cluster extent). The x- and y-axis indicate the time of testing and training sets after the stimulus onset, respectively. Red triangles indicate the mask onset time (60 ms after stimulus onset). The orange and green line plots under each TG map show the dynamic and stable index summarizing the temporal coding of TG, respectively. The x-axis shows the time after stimulus onset. The y-axis shows the magnitude of stable and dynamic index. Shaded areas represent ±1 bootstrapped standard error. CI: Neural code index.

Next, we performed a temporal generalization (TG) analysis^19^ on these four datasets in order to investigate the spatiotemporal neural dynamics underlying perceptual integration as a function of conscious access. Specifically, we trained the Kanizsa vs. control classifier at a particular time point and then generalized it to EEG signals over the entire period. We iterated this procedure until the Kanizsa vs. control classifier trained at every time point was generalized to the EEG signals obtained at every time point, to make a two-dimensional TG matrix of decoding accuracies (10-fold cross-validation). We computed the decoding accuracy at every time point (in 2 ms steps) using a sliding 20 ms window. We performed the TG analysis separately for the three electrode clusters (37 posterior, 37 central, and 36 anterior electrodes) to examine the role of each region in perceptual integration (Three rows of two-dimensional TG maps in Fig 2b).

As TG methods reveal whether the neural code that supports above-chance decoding of integrated percepts is stable or dynamically evolving over time, we also summarized these patterns of two-dimensional modulations of decoding accuracies in the TG map using a time-resolved dynamic and stable index^20^ (Line plots below each TG map in Fig 2b. See Methods for the details). Stable neural code refers to a situation where the Kanizsa vs. control classifier trained at a particular time point can reflect the presence of integrated percepts over a different period, resulting in the rectangle-shaped decoding accuracy modulations in the TG map. Dynamic neural code refers to a situation where the Kanizsa vs. control classifier trained at a particular time point can reflect the presence of integrated percepts only at the matched time point, resulting in the significant decoding accuracy modulations along the diagonal axis in the TG map. Briefly, the dynamic index is the proportion of the off-diagonal elements that were significantly smaller than the corresponding on-diagonal elements. The stable index is the proportion of the off-diagonal elements that were significantly greater than 0.5 and not significantly smaller than the corresponding on-diagonal elements.

In Figure 2b, the four columns, from left to right, represent Dataset 1 through Dataset 4 and the three rows from top to bottom represent the TG matrices and their time-resolved dynamic and stable index for the posterior, central and anterior regions, respectively. This table summarizes how the presence of integrated percepts is reflected in different brain regions over time as conscious access to the identical stimulus varied. Our first overall observation of the TG maps was that in most of the datasets except Dataset 4, the TG maps for all three brain regions showed significant time clusters where Kanizsa vs. control classification accuracy was significantly higher than chance, as indicated by the black contours. The temporal neural codes summarized by dynamic/stable index showed that both the posterior and central regions initially used dynamic neural code and later switched to stable neural code (The 1^st^ and 2^nd^ row in Fig 2b), while stable neural code was dominant in the anterior region (The 3^rd^ row in Fig 2b).

When a Kanizsa triangle was detected with a clear sense of its orientation, all three brain regions almost concurrently showed the significant decoding accuracy modulations in the TG maps (The 1^st^ column in Fig 2b), which were effectively captured by stable neural codes during a period from ~400 to 1000 ms after the stimulus onset (Green line plot under each TG map in the 1^st^ column of Fig 2b). Dynamic neural code, in contrast, briefly appeared from ~100 ms and ~200 ms after the stimulus onset in the posterior and the central region, respectively, whereas dynamic neural code was nearly absent in the anterior region (Orange line plot under each TG map in the 1^st^ column of Fig 2b). When the stimulus awareness was minimal, significant time cluster was observed only in the anterior area during a period from ~430 ms after the stimulus onset (The 4^th^ column in Fig 2b). Dataset 2 and Dataset 3 showed distinctly different spatiotemporal dynamics, despite their similar subjective Kanizsa visibility ratings, slightly above and below the grand average Kanizsa visibility rating of 3.5. That is, significant time cluster in the TG map appeared increasingly later along the postero-anterior direction after stimulus onset for Dataset 2 (The 2^nd^ column in Fig 2b), whereas for Dataset 3, the anterior region showed significant time cluster earlier than the posterior and central regions (The 3^rd^ column in Fig 2b). This reversal of temporal dynamics across brain regions was shown by time-resolved stable indices for the three brain regions. In Dataset 2, stable neural code appeared from ~480 ms in the posterior and central regions while the anterior region showed stable neural code from ~560 ms after the stimulus onset (Green line plot under each TG map in the 2^nd^ column of Fig 2b). However, in Dataset 3, the anterior, central and posterior region showed stable neural codes starting at ~300 ms, ~400 ms, and ~500 ms, respectively (Green line plot under each TG map in the 3^rd^ column of Fig 2b). The temporal dynamics of dynamic neural codes in each brain area were not significantly different from those shown in Dataset 1 (Orange line plot under each TG map in the 2^nd^ & 3^rd^ columns of Fig 2b). These results suggest that spatiotemporal neural dynamics underlying perceptual integration change depending on the degree of conscious access to the same stimulus. Our TG results do not significantly change when we divided trials into five datasets that exhaustively used all trials (Fig S1).

Next, we examined how dynamic/stable index varied as a function of conscious access in each brain region. In the posterior region, we observed increasingly delayed dynamic index peak times of ~265 ms, ~427 ms, and ~500 ms after the stimulus onset as conscious access decreased from Dataset 1 to Dataset 3 (Orange line plots in the 1^st^ row in Fig 2b). In contrast, stable indices became dominant during a very late period from ~600 ms to 1000 ms after the stimulus onset in Dataset 1-3 (Green line plots in the 1^st^ row in Fig 2b). The central region did not show any noticeable systematic change in dynamic and stable index from Dataset 1 to Dataset 4 (Orange and Green line plots in the 2^nd^ row in Fig 2b). The anterior region showed the most striking change in stable index as a function of conscious access. When the Kanizsa visibility decreased from Dataset 1 to Dataset 2, the time for the stable index to appear was delayed from ~420 ms to ~560 ms after the stimulus onset (Green line plots in the 1^st^ and 2^nd^ column, the 3^rd^ row in Fig 2b). However, when the target identification performance was at chance level and the average Kanizsa visibility was just below 3.5 (Dataset 3), the anterior region showed significant decoding accuracies over the off-diagonal areas starting from ~300 ms after the stimulus onset (Green line plot in the 3^rd^ column, 3^rd^ row in Fig 2b). This stable neural code in the anterior region remained even when the stimulus awareness was minimal (Dataset 4) unlike the posterior and central regions. This non-monotonic decoding accuracy modulation in the TG map of the anterior region as a function of conscious access strongly suggests the role of the anterior region in perceptual integration at low Kanizsa visibility.

To examine if the discrimination between the four Kanizsa stimuli and control stimulus was affected by a specific Kanizsa stimulus type, we trained and tested a decoder to classify the four Kanizsa stimuli (Fig S2). This analysis showed that neural responses to the four Kanizsa stimuli were not significantly different from each other. We then performed a decoding analysis to classify the trials of each Kanizsa stimulus against the control trials (Fig S3). Given that all four types of Kanizsa trials were robustly classified against the control trials, any potential biases introduced by specific Kanizsa types are likely to be minimal. We also investigated if the neural response to non-Kanizsa control stimulus was influenced by subjective (imagined) conscious access to Kanizsa figure or not (Fig S4). In the same way we sorted Kanizsa trials in Figure 2, we sorted all 200 non-Kanizsa control trials in order from correct trials ranked by visibility rating to incorrect trials ranked by visibility rating. Then we divided these control trials into two groups of 100 trials. Our method did not reliably classify the higher versus lower subjective conscious access of Kanizsa figures on the non-Kanizsa control trials which did not induce perceptual integration. These results suggest that decoding accuracy modulations over the TG maps primarily reflect subjective visibility of Kanizsa figures associated with perceptual integration.

## Discussion

Our goal was to characterize the spatiotemporal neural dynamics underlying perceptual integration under different levels of conscious access. To this end, we applied the temporal generalization (TG) method to scalp-recorded EEG to reveal how integrated percepts of Kanizsa figures are stably and/or dynamically decoded in the anterior, central, and posterior regions, using near-threshold stimuli that generated various degrees of visibility from trial to trial. We found that with decreasing conscious access, the posterior region showed dynamic-to-stable neural-code transition that occurred increasingly later after the stimulus onset and eventually disappeared (The 1^st^ row in Fig 2b). However, the anterior region primarily showed stable neural code (The 3^rd^ row in Fig 2b) that occurred increasingly later with decreasing conscious access. However, when behavioral performance was at chance level and Kanizsa visibility was just below the grand average visibility rating of 3.5, the stable neural code appeared much earlier than when Kanizsa stimuli were more visible. This stable neural code in the anterior region still remained even when Kanizsa visibility was minimal. These findings reveal distinct brain mechanisms underlying perceptual integration depending on the degree of conscious access.

### Spatiotemporal dynamics of perceptual integration are different when stimulus awareness is maximal versus minimal

Our results have a number of implications for understanding neural processes underlying seen and unseen sensory information. First, our findings provide new insights into how the brain processes perceptual integration of maximally and minimally visible stimuli when the stimulus awareness was influenced only by internal states. When trials were sorted according to the degree of conscious access, that is, subjective visibility rating and behavioral accuracy, the spatiotemporal neural dynamics supporting the decoding of perceptual integration of maximally and minimally visible stimuli were clearly distinct from each other even though the physical intensity of the stimuli was identical (compare the 1^st^ and 4^th^ column in Fig 2b).

At maximal stimulus awareness, the observed transition from dynamic to stable neural code in the posterior and central region and the dominant stable neural code in the anterior region were quite different from the robust square-shaped decoding of categories and exemplars in occipital, ventral-temporal, parietal and prefrontal areas in a recent electrocorticography (ECoG) study^23^. This difference in the shape of the significant decoding is likely due to a difference in stimulus awareness; whereas the ECoG study used clearly visible stimuli presented for 300 ~ 1500 ms, we used briefly presented backward-masked stimuli. This suggests that both the brief stimulus presentation and the backward masking change square-shaped stable decoding dynamics into the observed temporal dynamics, even though our stimulus was subjectively highly visible in Dataset 1.

When the physically identical stimuli were subjectively perceived as nearly invisible, the stable neural code appeared from ~400 ms after the stimulus onset only in the anterior region (The 4^th^ column in Fig 2b). In contrast, at maximal stimulus awareness, all three brain regions almost concurrently showed the stable neural code during a period from ~400 to 1000 ms after the stimulus onset (Green line plots in the 1^st^ column in Fig 2b).

### Frontal areas are differentially involved in processing perceptual integration depending on subjective visibility

We observed distinctive changes in spatiotemporal neural dynamics around the average visibility rating of 3.5. The TG map for the anterior region showed that the cluster of time points of significant decoding of integrated percepts increased as the visibility rating decreased from 4.0 to 3.0 (The 3^rd^ row of Fig 2b and Fig 3b). This is likely due to the involvement of frontal areas in rescuing information that was compromised at chance level of behavioral performance. A previous behavioral study reported that postcued attention could reach into the past and bring a stimulus too faint to see into consciousness by acting on the memory trace of the stimulus that disappeared before being attended^24^. Consistent with recent research suggesting that such temporally flexible conscious perception can be implemented through long-distance information sharing^25–28^, our results suggest that perceptual integration was processed earlier in sustained activity patterns in the anterior region than in the posterior and central regions at chance level of behavioral performance (The 3^rd^ column of Fig 2b).

**Figure 3.**
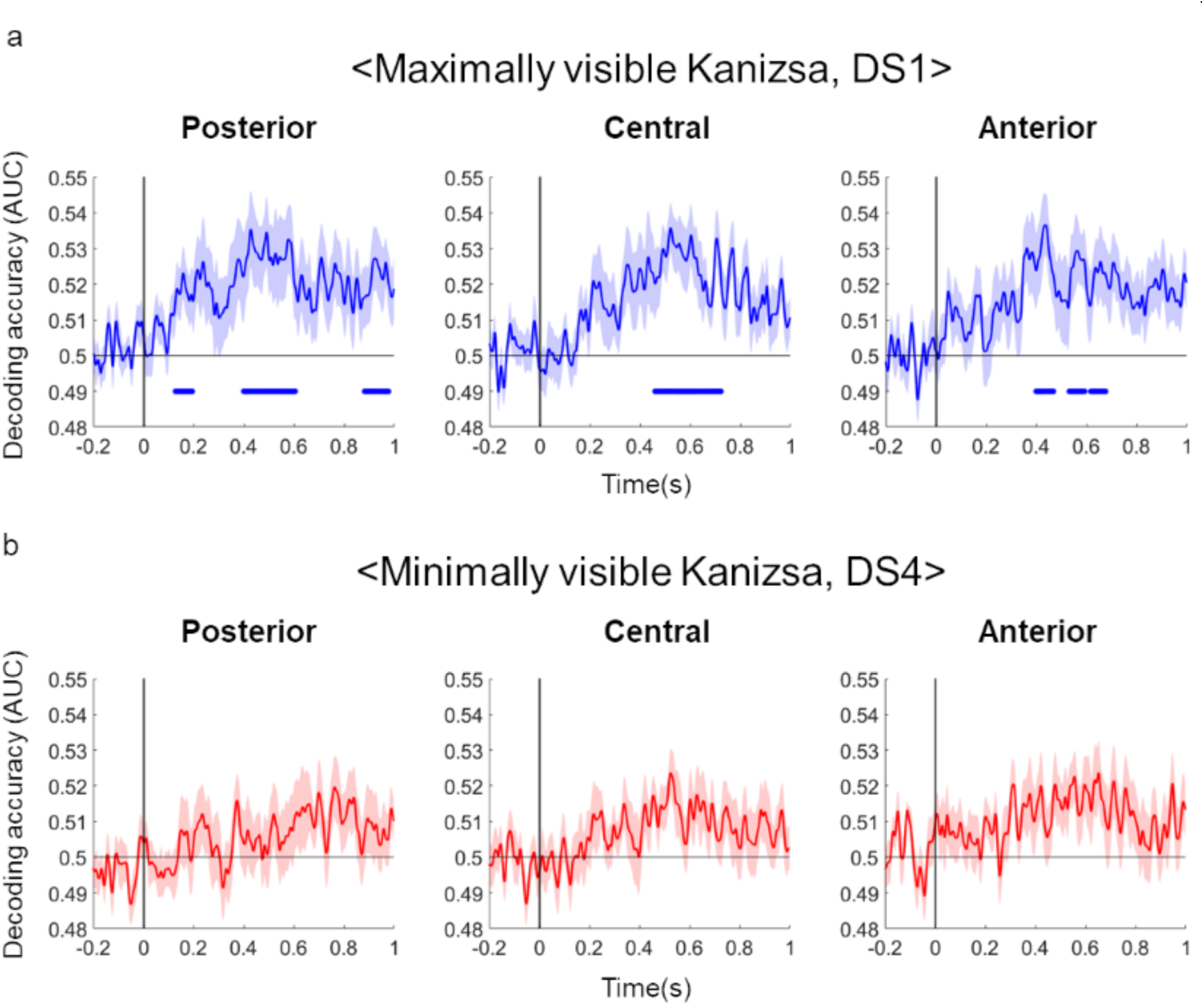
Diagonal part of TG map. **a.** The Kanizsa vs. control classification accuracy as a function of time for maximally visible Kanizsa stimuli (Dataset 1) for three brain regions. Blue graph in each column indicates the diagonal part of the TG map in each brain region (the 1^st^ column of Fig 2b). Horizontal blue lines below each graph represent statistically significant time points of above-chance decoding accuracy (> 0.5) (*p* < 0.05, based on cluster extent). **b.** The Kanizsa vs. control classification accuracy as a function of time for minimally visible Kanizsa stimuli (Dataset 4) for three brain regions. Red graph in each column indicates the diagonal part of the TG map in each brain region (the 4^th^ column of Fig 2b). The x-axis shows the time after stimulus onset and the y-axis shows the decoding accuracy (AUC). Shaded areas represent ±1 SEM.

Frontal areas are likely to play a crucial role in retrieving stimuli, especially near threshold. The TG maps for the anterior region revealed that the significant cluster was the largest at the visibility rating of 3.0 and nearly disappeared at the rating of 1.6, exhibiting a sharp decline between 3.0 and 1.6 (Figs 2b). These findings are reminiscent of a previous source-imaged EEG study demonstrating that diagonal-shaped stable decoding of auditory sound became robust when the signal-to-noise ratio of auditory sound was changed from below-threshold to threshold audibility level^16^. In our case, however, the distinctive changes in stable decoding dynamics are purely due to internal variations in conscious access, not confounded by physical stimulus intensity changes as in the previous study. Although this same study demonstrated the late sustained frontal activations during conscious processing in the absence of report^16^, we do not exclude the possibility that the late stable neural code in the anterior region is due to task-related reports at the end of the trial. However, the earlier appearance of the stable neural code in the anterior region than in the central and posterior regions (The 3^rd^ column of Fig 2b) suggests that frontal regions are involved in conscious perception of inconspicuous stimuli. The residual significant decoding during a period from 400 – 1000 ms in the frontal region without any significant decoding in the posterior and central regions at the minimal stimulus awareness condition (The 4^th^ column of Fig 2b) suggests that the frontal region attempts to rescue nearly invisible stimulus but fails broadcast the recovered information to a global network of areas.

In contrast, when the visibility rating was above the average visibility rating of 3.5, the simultaneous or earlier appearance of sustained activations in the posterior region than in the central and anterior regions (Figs 2b) suggests the major role of the posterior region in processing conspicuous stimuli. These results strongly indicate that brain implements differential mechanism underlying perceptual integration depending on the degree of the stimulus awareness, which provides interesting clues about the role of the frontal lobes in conscious perception, which is currently hotly debated^29,30^.

### Conscious access is variable around the chance level of behavioral accuracy

The observed subjective-visibility dependent spatiotemporal neural dynamics underlying perceptual integration furthers our understanding of the relation between perceptual integration and conscious access. Previous studies using the attentional blink paradigm or the masked priming paradigm without reports of the target visibility have traditionally assumed that there is no conscious access at all in the condition where participants performed at the chance level of behavioral accuracy^4,5,7–9^. Based on this tacit assumption, these previous studies have claimed that there is a clear dissociation between conscious access and perceptual or semantic integration. However, our results show that for chance-level behavioral performance, there is a mix of slightly visible and invisible trials around the average visibility rating point. This is consistent with a previous study demonstrating the non-monotonic profile of the standard deviation of audibility with increasing SNR, with a burst in variability around detection threshold^16^. This indicates that conscious access to stimuli above or below threshold may be an all-or-none phenomenon^31^, but conscious access to the threshold-level stimulus is a continuous rather than an all-or-none phenomenon. For this reason, one should be careful to use the terms such as “unconscious” or “without conscious access” when describing the stimulus awareness at the chance-level behavioral performance on the threshold stimulus.

### Neural representations of perceptual integration are temporally distributed

The diagonal-shaped stable decoding appeared in all three brain regions across subjective visibility levels except for the minimal stimulus awareness (Fig 2b). That is, the classifier discriminating Kanizsa illusion at a particular time point could decode the presence of integrated percepts stably over a period of a few hundred milliseconds in each brain region, resulting in the rectangular shaped modulations in the TG map along the diagonal. A recent attentional blink study demonstrated that the second target Kanizsa vs. control classification accuracy was well above chance around 200 ms in the posterior region by applying Kanizsa vs. control classifier trained at a particular time point to only the matched time point^5^. When the second target Kanizsa was followed by a strong high-contrast mask, the behavioral accuracy and the decoding accuracy were completely wiped out. Based on those observations, authors have suggested that dynamically changing activity patterns in the posterior region underlie the presence of integrated percepts. Consistent with this previous study, when we extracted the diagonal part of the TG map in the posterior region (The 1^st^ row in Fig 2b), the decoding accuracy for maximally visible Kanizsa stimuli was significant around 200 ms after Kanizsa stimulus onset (The 1^st^ column in Fig 3a) and was around chance level for minimally visible Kanizsa stimulus (The 1^st^ column in Fig 3b). For maximally visible Kanizsa stimuli in the current study, we additionally observed the significant decoding accuracy after 400 ms post-stimulus onset in all three regions (Fig 3a). However, these results are part of a TG map that showed the significant off-diagonal decoding performance in all three brain regions, so they provide only a partial view of the brain dynamics underlying perceptual integration. Thus, the TG method demonstrates a full view of the broadly distributed representation of perceptual integration by complementing a previous study that examined only the diagonal part of the TG map of perceptual integration in posterior regions^5^.

Moreover, significant decoding of the presence of integrated percepts across three brain regions with differential temporal dynamics (The first three columns in Fig 2b) are reminiscent of the Global Neuronal Workspace Theory (GNWT)^32,33^ predicting that conscious access is associated with qualitative changes in the configuration of activity throughout the brain^33–35^. The observed subjective visibility-dependent differential spatiotemporal neural dynamics seem to be consistent with GNWT positing that conscious access arises through the temporally flexible broadcasting of sensory information to a global network of areas^26^. Especially when the stimulus becomes less visible, the past sensory information may be reactivated within anterior region, amplified and broadcast within a global workspace (The 3^rd^ column of Fig 2b), leading to the temporal flexibility of conscious access. However, at least for Kanizsa illusion seen above the average visibility rating of 3.5, we cannot rule out the possibility that the observed dominant diagonal-shaped significant decoding in the posterior region lasting for a sustained period from 200 ms to 1000 ms from the target onset is consistent with the prediction of the Integrated Information Theory (IIT). According to IIT, sustained activity in the posterior “hot zone” increases and decreases in correspondence with conscious sensory experiences^22^. Taken together, while further investigation is needed to determine which of the conflicting theories of consciousness is valid, our findings suggest that the neural mechanisms underlying conscious access may vary with subjective visibility.

In conclusion, our study examined the neural representations of perceptual integration under different levels of conscious access to stimulus information using constant near-threshold stimuli whose visibility spontaneously varied. We found that the temporal dynamics of the broadly distributed representation of perceptual integration varied with subjective visibility. Our results contribute to the current theories of consciousness by shedding light on the relationship between conscious access and perceptual representation.

## Materials and Methods

### Participants

Twenty-four observers participated in the experiment. Four observers whose EEG data were excessively noisy due to bad recording preparation, resulting in high artifacts including eye and body movement, were excluded from the analysis (n=20, mean age 23.6 ± 2.15, eleven females, nine males). All observers had normal or corrected-to-normal vision, gave written consent to participate as paid volunteers, and were tested in a dark room. All observers were naïve to the purpose of the experiment. The study was approved by the Institutional Review Board of the Korea National Institute for Bioethics Policy.

### Stimuli

All visual and auditory stimuli were generated and presented using Psychophysics Toolbox 6– 11 along with custom scripts written for MATLAB (Mathworks Inc.) running on a Mac mini. The 22” CRT (ViewSonic PF817) monitor was set to a 100 Hz refresh rate and a resolution of 800 × 600 pixels. Both temporal and luminance calibrations were performed using a calibrated photocell, and monitor gamma tables were adjusted to ensure response linearity and a constant mean luminance of 22 cd/m². The observers’ viewing distance from the monitor was 60 cm.

We constructed a set of four Kanizsa triangles and one control stimulus (Fig 1). All stimuli were made with four pacmans. Each trial consisted of a target presentation followed by backward masking. Each trial was preceded and followed by a blank period. All target and mask stimuli consisted of an identical set of four pacmans (contrast, 80%; visual angle, 5.3°; color, black; background color, gray), and were distinguished only by the relative orientations of the four pacmans.

### Experimental Procedure

The observer initiated each trial by pressing the space bar. A central fixation cue appeared for 500ms. Subsequently, they viewed either a Kanizsa or a control target which was followed by a mask (Fig 1a). Targets were presented in a random sequence and could be either a Kanizsa or a control stimulus. Each target was presented for 20ms, followed by a 20ms mask presented after a 40ms blank interval. During this entire period, observers were instructed to maintain fixation on the center of the screen and attempt to withhold eyeblinks. After the mask, a central fixation cue appeared for 920ms. The instructions for task responses were then displayed. Observers pressed a number key to indicate the target stimulus that they thought they saw (Kanizsa triangle pointing upper right, Kanizsa triangle pointing upper left, Kanizsa triangle pointing lower left, Kanizsa triangle pointing lower right, or the control stimulus [no Kanizsa triangle]), and then rated the visibility of a Kanizsa triangle (0 to 7). The visibility-rating scheme was based on previous studies^31,36^. Specifically, observers were instructed to respond 0 when they did not see a Kanizsa triangle at all, respond 1-3 when they had a vague impression of a Kanizsa triangle depending on the precision of this impression, respond 4-6 when they have seen a Kanizsa triangle depending on the clarity and precision of their target perception, and respond 7 when they saw the clearest possible Kanzsa triangle under the brief and masked presentation. Each block consisted of 100 trials (including 20 trials for each of the five target stimuli), and a total of 10 blocks were performed.

### EEG signal acquisition and preprocessing

EEG data were recorded with 128-sensor HydroCel Sensor Nets (Electrical Geodesics, Eugene OR) at a sampling rate of 500 Hz. Offline preprocessing and analysis were performed using Fieldtrip^37^, EEGLAB^38^, and custom scripts in MATLAB (Mathworks Inc.). EEG data were band-pass filtered from 0.3 to 200 Hz. We used Artifact Subspace Reconstruction (ASR) routine (Mullen et al., 2015) to remove noisy channels, and re-referenced all data using the average as the reference (Bidgely-Shalmo et al., 2015). We removed line noise (60, 120, and 180 Hz) using the *cleanline* EEGLAB plugin. Then, the artifactual response components induced by eye movements and blinks were identified vis independent component analysis (ICA) (Makeig et al., 1996). Finally, we determined ICs that were classified as artifacts using *ADJUST* EEGLAB plugin (Mognon et al., 2011), and removed them from the data. The EEG data were epoched from −500ms to 1000ms relative to the onset of the target stimulus and baseline-corrected by subtracting the average signal from −200 to −50ms. All analyses were performed with 111 EEG channels, excluding 17 channels vulnerable to movement artifacts including electrodes around the ears and on the face.

### Temporal generalization (TG) method

To characterize the neural dynamics of perceptual integration, we separately employed the Linear Discriminant Analysis (LDA) on each of the four datasets at one time point. We then applied the obtained weights to the test set at all time points, using a sliding window of 20 ms with 2 ms steps. This procedure was repeated, allowing us to calculate the area under the curve (AUC) for each time point and create a two-dimensional temporal generalization matrix of AUC. The AUC was computed based on the true positive rate (TPR) and false positive rate (FPR). For most results, we performed 10-fold cross-validation (CV). To assess the significance of the AUC values being greater than 0.5, we used non-parametric cluster-based permutation testing (1000 permutations) using one-sample *t*-tests^39^. We did not have separate training dataset that had no backward mask stimulus and no behavioral report. But it has been shown that switching to no-report makes the visibility of the stimuli different^40^. This was one reason why we directly used our main experimental data as training and test datasets.

Additionally, we extracted the diagonal part from the TG results (Fig 3), specifically focusing on the data corresponding to the same time points of training and testing. For these comparisons, we employed non-parametric cluster-based permutation testing with 200 permutations and conducted two-sample *t*-tests to evaluate whether the AUC was greater than 0.5.

Three electrode clusters (37 posterior-central, 37 central, and 36 anterior-central) were used in Fig 2b.

### Data sorting procedure for the TG analysis

We sorted Kanizsa trials in order from correct trials ranked by visibility rating to incorrect trials ranked by visibility rating. We repeated the same trial sorting procedure for each Kanizsa stimulus type. From these sorted trials, we selected the top 50 and bottom 50 trials of each of the four Kanizsa triangle types to create Dataset 1 (200 trials in total) and Dataset 4 (200 trials in total), respectively. Next, we created Dataset 3 by extracting trials in which behavioral performance was around chance level of 0.2. As the average visibility rating of Dataset 3 was 3 which was just below the grand average visibility rating of 3.5, we created Dataset 2 by extracting trials that made the average visibility rating higher than the grand average visibility rating by 0.5. In this way, Dataset 2 and Dataset 3 mainly consisted of trials with ambiguous conscious access, such as correct trials with low visibility ratings or incorrect trials with high visibility ratings, and they were positioned between Dataset 1 and Dataset 4. Thus, trials were split into four datasets of equal number of trials: the maximal visibility dataset (Dataset 1, visibility *M* = 5.3, performance correct *M* =0.98), above-mean visibility dataset (Dataset 2, visibility *M* = 4, performance correct *M* =0.60), below-mean visibility dataset (Dataset 3, visibility *M* = 3, performance correct *M* =0.16), and minimal visibility dataset (Dataset 4, visibility *M* = 1.6, performance correct *M* =0.08).

Because the four datasets did not use all Kanizsa trials for the TG analysis shown in Fig 2, we conducted an additional TG analysis using all Kanizsa trials (Fig S1). As with the analysis shown in Fig 2, we sorted Kanizsa trials in order from correct trials ranked by visibility rating to incorrect trials ranked by visibility rating. All Kanizsa trials were divided into the five datasets of equal number of trials. Each of the five datasets had 11~12 overlapping trials. For example, if there were 200 trials for each of the four Kanizsa, trials 1-50 were extracted for dataset 1, trials 38-87 for dataset 2, trials 75-124 for dataset 3, trials 113-162 for dataset 4, and trials 151-200 for dataset 5 from the one Kanizsa. This process was repeated for all four Kanizsa, resulting in a total of 200 trials for each dataset.

### Dynamic/stable index

To quantify the magnitude of dynamic and stable coding, we calculated a dynamic/stable index^20^. For the dynamic coding, we tested whether the off-diagonal element of the matrix TG(t1,t2) was significantly smaller than the corresponding on-diagonal elements of the matrix TG(t1,t1) and TG(t2,t2)

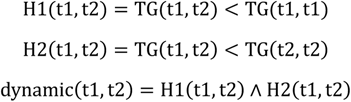

We used 40 ms square window for each time point.

For the stable coding, we tested whether the off-diagonal element of the matrix TG(t1,t2) was significantly higher than zero and, at the same time, not significantly smaller than the corresponding on-diagonal elements of the matrix TG(t1,t1) and TG(t2,t2):

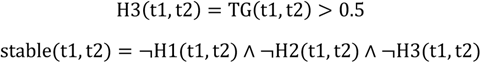

We gained the indices of stable or dynamic coding at a time point t by calculating the proportion of significant stable or dynamic coding elements over a 40ms square window centered around time point *t*. We simultaneously controlled the temporal smearing effect caused by a 40ms moving-average filter by excluding the elements within ±20ms of the diagonal axis from the analysis. An index value (computed separately for dynamic and stable coding) of 1 indicates that the TG is completely dynamic or stable, 0 indicates that the TG is not at all dynamic or stable.

## Acknowledgment

This research was supported by IBS-R001-D2.

## Supplementary Figures

**Figure S1.**
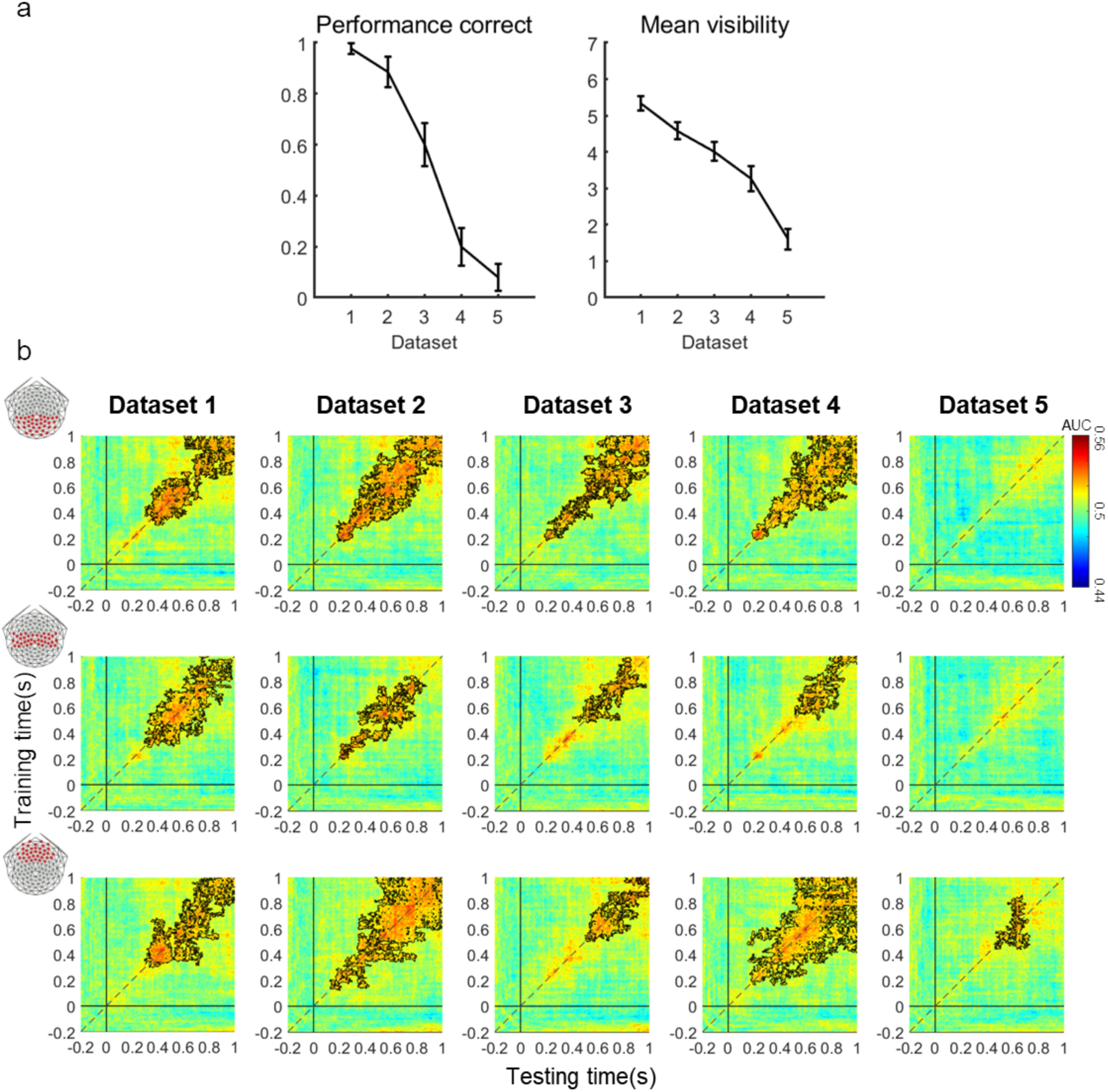
Spatiotemporal neural dynamics of perceptual integration do not substantially change when we divided trials into five datasets that used all trial. As with the analysis shown in Fig. 2, we sorted Kanizsa trials in order from correct trials ranked by visibility rating to incorrect trials ranked by visibility rating. All trials were divided into the five datasets of equal number of trials. Each of the five datasets had 11~12 overlapping trials. For example, if there were 200 trials for each of the four Kanizsa stimuli, trials 1-50 were extracted for dataset 1, trials 38-87 for dataset 2, trials 75-124 for dataset 3, trials 113-162 for dataset 4, and trials 151-200 for dataset 5. This process was repeated for all four Kanizsa stimuli, resulting in a total of 200 trials for each dataset. Here, Dataset 1, 3, and 5 correspond to Dataset 1, 2 and 4 in Fig 2b. **a.** Performance correct and average Kanizsa visibility rating as a function of dataset. **b.** TG maps of each dataset for the posterior, central, and anterior regions. Colors represent the decoding accuracy (AUC). Black contours indicate clusters of significant decoding accuracies above chance (p < 0.05, based on cluster extent). The x- and y-axis indicate the time of testing and training sets after the stimulus onset, respectively.

**Figure S2.**
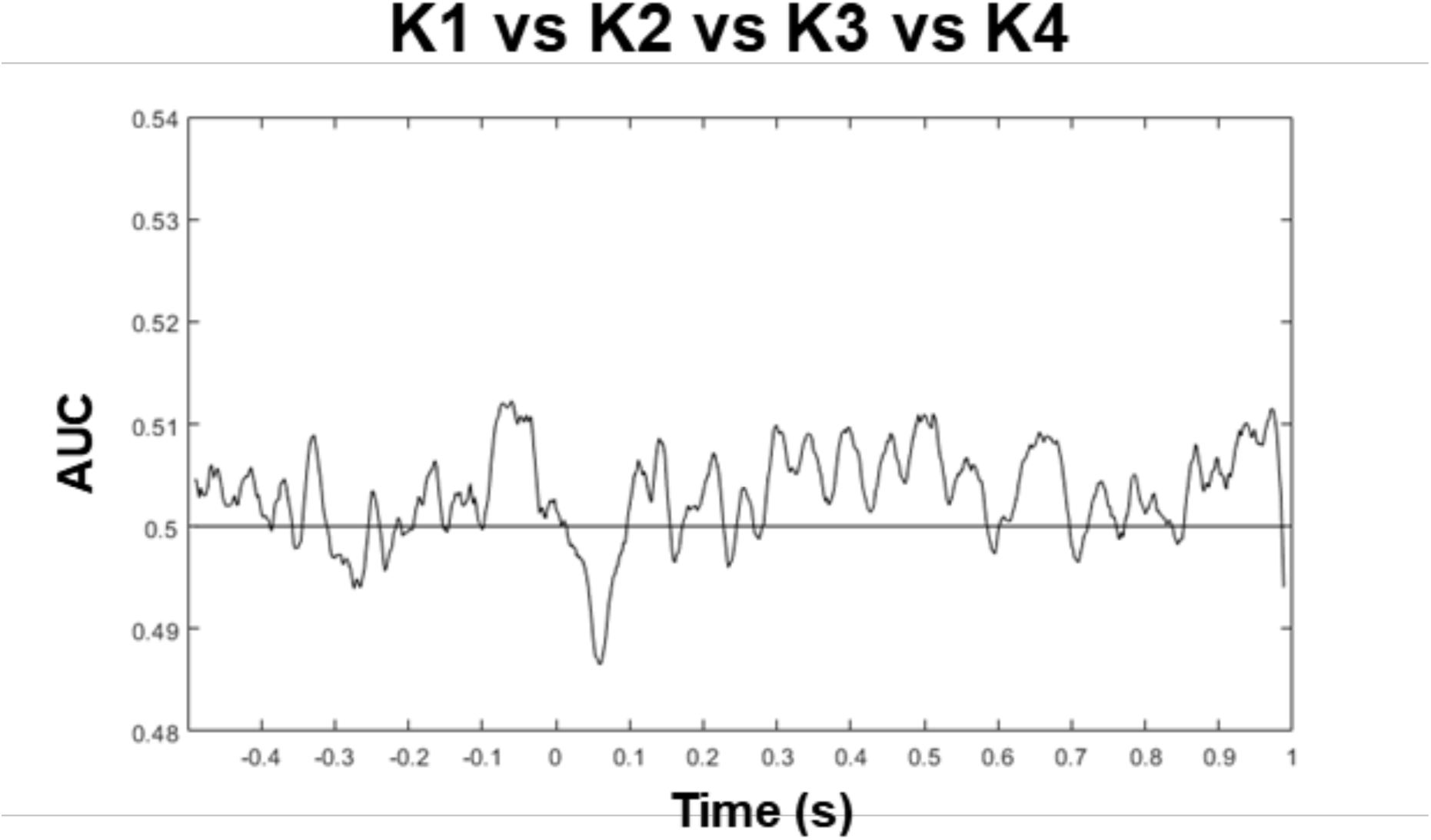
Multivariate neural responses to the four types of Kanizsa stimuli are not different from each other. The decoder that was trained to classify the four types of Kanizsa stimuli at a particular time point was applied to the matched time point (10-fold cross validation). The x-axis shows the time after stimulus onset and the y-axis shows decoding accuracy (AUC). K1~K4: The four types of Kanizsa stimuli are shown in Fig 1.

**Figure S3.**
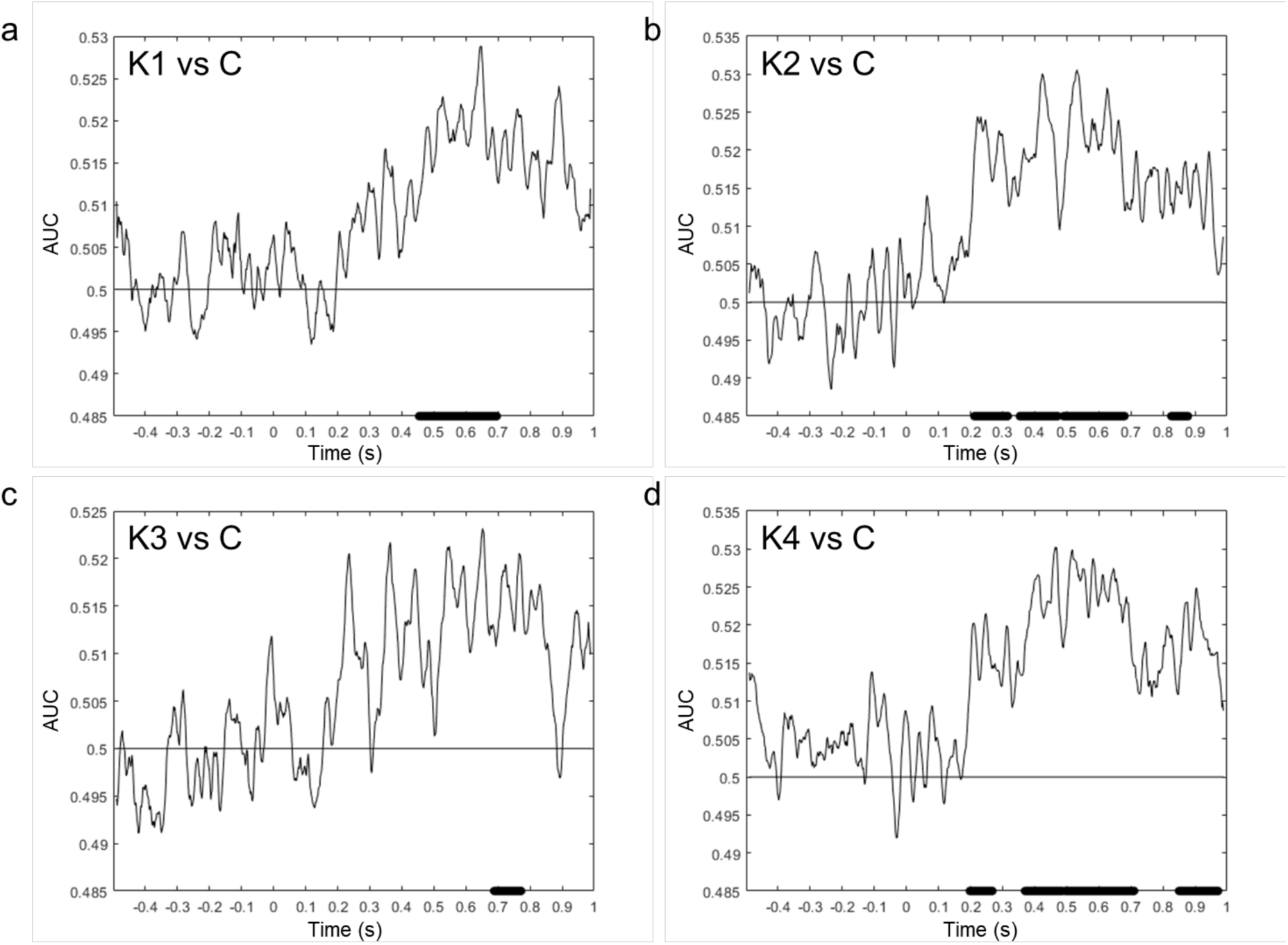
Multivariate neural responses discriminate each type of Kanizsa stimuli from non-Kanizsa control stimulus. The Kanizsa stimulus type vs. control decoder trained at a particular time point was applied to the matched time point (10-fold cross validation). The x-axis shows the time after stimulus onset and the y-axis shows the decoding accuracy (AUC). Horizontal black bars on the x-axis indicate the time points of significant decoding of each Kanizsa stimulus type (p < 0.05, based on cluster extent). K1~K4 and C: The four possible types of Kanizsa stimuli and non-Kanizsa control stimulus are shown in Fig 1.

**Figure S4.**
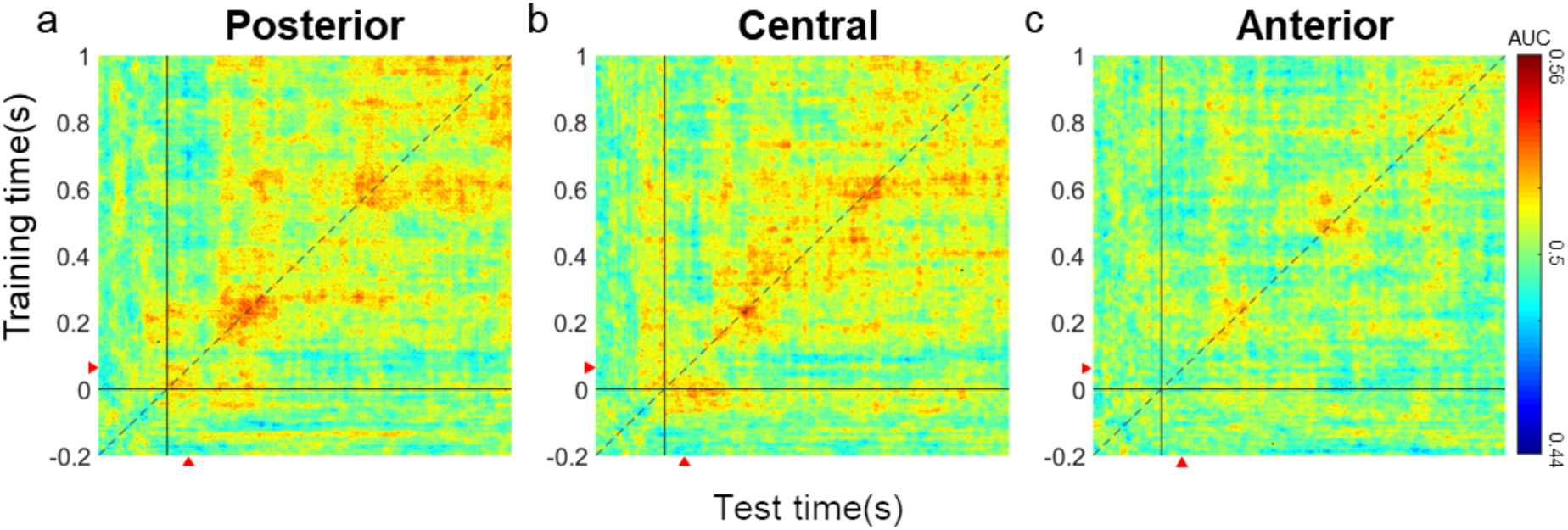
Multivariate neural responses do not distinguish between the split-half high-visibility and low-visibility non-Kanizsa control trials. **a-c.** In the same way we sorted Kanizsa trials in Figure 2, we sorted all 200 non-Kanizsa control trials in order from correct trials ranked by visibility rating to incorrect trials ranked by visibility rating. Then we divided these control trials into two groups of 100 trials. The decoder that was trained to discriminate these two groups of non-Kanizsa control trials at a specific time point was tested across all time points in three brain regions shown in two-dimensional decoding accuracy (AUC) modulations in the TG map. Colors indicate the decoding accuracy (AUC) at each pair of train-test time. There was no significant time cluster where the decoding accuracies were greater than 0.5 (*p* = 0.05, based on cluster extent). The x- and y-axis show the train and the test time after control stimulus onset, respectively. Red triangles indicate the time of mask onset.

